# Multiple founding paternal lineages inferred from the newly-developed SNPSeqTyper 639 Y-SNP panel suggested the complex admixture and migration history of Chinese people

**DOI:** 10.1101/2022.12.20.520342

**Authors:** Guanglin He, Mengge Wang, Jing Chen, Lei Miao, Jie Zhao, Qiuxia Sun, Shuhan Duan, Zhiyong Wang, Xiaofei Xu, Yuntao Sun, Yan Liu, Jing Liu, Zheng Wang, Lanhai Wei, Chao Liu, Jian Ye, Le Wang

## Abstract

Non-recombining regions of the Y-chromosome are inherited male-specifically and recorded the evolutionary traces of male human populations. Recent whole Y-chromosome sequencing studies have identified previously unrecognized population divergence, expansion and admixture processes, which promotes a better understanding and application of the observed patterns of Y-chromosome genetic diversity. Here, we developed one highest-resolution Y-SNP panel for forensic pedigree search and paternal biogeographical ancestry inference, which included 639 phylogenetically informative SNPs (Y-SNPs). We genotyped these loci in 1033 Chinese male individuals from 33 ethnolinguistically diverse populations and identified 257 terminal Y-chromosomal lineages with frequency ranging from 0.010 (singleton) to 0.0687. We identified six dominant common founding lineages associated with different ethnolinguistic backgrounds, which included O2a2b1a1a1a1a1a1a1-M6539, O2a1b1a1a1a1a1a1-F17, O2a2b1a1a1a1a1b1a1b-MF15397, O2a2b2a1b1-A16609, O1b1a1a1a1b2a1a1-F2517 and O2a2b1a1a1a1a1a1-F155. The AMOVA and nucleotide diversity estimates revealed considerable differences and high genetic diversity among ethnolinguistically different populations. We constructed one representative phylogenetic tree among 33 studied populations based on the haplogroup frequency spectrum and sequence variations. Clustering patterns in principal component analysis and multidimensional scaling results showed a genetic differentiation between Tai-Kadai-speaking Li, Mongolic-speaking Mongolian and other Sinitic-speaking Han Chinese populations. Phylogenetic topology inferred from the BEAST and Network relationships reconstructed from the popART further showed the founding lineages from culturally/linguistically diverse populations, such as C2a/C2b was dominant in Mongolian people and O1a/O1b was dominant in island Li people. We also identified many lineages shared by more than two ethnolinguistically different populations with a high proportion, suggesting their extensive admixture and migration history. Our findings indicated that our developed high-resolution Y-SNP panel included major dominant Y-lineages of Chinese populations from different ethnic groups and geographical regions, which can be used as the primary and powerful tool for forensic practice. We should emphasize the necessity and importance of whole-sequencing of more ethnolinguistically different populations, which can help identify more unrecognized population-specific variations for the final promotion of Y-chromosome-based forensic applications.

## INTRODUCTION

Human genomics studies in the whole-genome sequencing era have updated our understanding of the patterns of genetic diversity among ethnolinguistically diverse worldwide populations, fine-scale population structure, variation spectrum of various kinds of genetic variations of single nucleotide variations (SNV), structural variations (SV) and complex mobile elements [1-3]. The properties of the non-recombining region of the human Y-chromosome (NRY), that is, male specificity, haploidy, and absence of crossing over, make its genetic variations a powerful tool in evolutionary studies and forensic investigations, especially in cases where standard autosomal short tandem repeat (STR) profiling is not informative [4, 5]. Haplotypes composed of Y-STRs or Y-SNPs have been applied to characterize the paternal lineages of unknown male contributors or infer paternal biogeographical ancestry [5-7]. Y-SNPs define stable haplotypes as haplogroups, which could be used to construct robust phylogeny. A large body of research has been dedicated to identifying novel Y-chromosomal genetic variations, detailing the phylogenetic tree, and characterizing the patterns of differentiated paternal lineages [7-9]. Nowadays, the advances and applications of next-generation sequencing (NGS) technology provide unbiased ascertainment of Y-SNPs, leading to the construction of detailed phylogenies in which branch lengths are proportional to divergence times and enabling the estimates of time to the most recent common ancestor (TMRCA) of branch nodes [7, 10, 11]. The Y-chromosomal molecular clock built a link between genetic diversity and human migration and admixture history, which could be used to estimate the time when a lineage originated or expanded or when an ancestral population split into two paternal populations and migrated into different areas.

Y-chromosome-based phylogeny was used to explore the population origin, admixture and evolutionary history at the era of the first genome sequencing era. Zerjal et al. genotyped over 32 markers among 2123 males and identified the genetic legacy of the Mongol western Expansion [12]. Similar studies focused on large-scale population cohorts in Mongolia and China revealed that regional population migration and admixture models contributed to the observed patterns of genetic diversity rather than simple cultural diffusion [13, 14]. The significant changes in the research patterns occurred when Wei et al. introduced the high-coverage complete Y-sequences in identifying the novel phylogenic variations and calibrating the Y-chromosomal phylogeny [15]. They reported 6662 high-confidence lineage informative SNPs (LISNPs) by analyzing 8.97 Mb of the NRY regions in 36 diverse genomes. Followingly, Karmin et al. sequenced 456 geographically diverse individuals and found that global cultural changes were associated with the identified bottleneck of Y-chromosome diversity based on the phylogenetic analysis of the high-coverage Y-chromosome sequences [16]. Poznik et al. focused on the 1244 whole Y-chromosome genomes from the 1000 Genomes Project and observed punctuated bursts in human male demography inferred from the identified ∼6000 variants [7]. Large-scale Y-chromosome phylogeny reconstruction from the European, Oceanian and Siberian populations further reported complex population origin tracts, gene flow events and population standstill, rapid population divergence and expansion [17-20]. The single population-scale Y-chromosome investigations focused on Chinese Tibetan, Han, Mongolian and other populations have been conducted, which have identified the population-specific founding lineages and corresponding population origin, separation, and following admixture events [10, 21-23]. However, large-scale Y-chromosome surveys based on high-resolution systems or whole-genome sequencing remained to be conducted to present the entire landscape of genetic diversity and fine-scale paternal genetic history.

The deeper understating of Y-chromosome variations in human evolutionary research further promoted its wide applications in forensic science [4, 24]. The Y-chromosomal phylogeny has become an essential pillar of forensic pedigree searches and paternal ancestry inference. On this basis, several Y-SNP panels of different resolutions have been developed and validated in geographically and linguistically diverse populations [25-31]. Due to the popularity of the capillary electrophoresis (CE) platform, currently available dedicated Y-SNP genotyping tools developed based on this system have restrictions on the number of Y-SNPs analyzed simultaneously [26, 28, 29, 32, 33], hindering the dissection of paternal biogeographical ancestry at high resolution. Here, targeted NGS technologies are up-and-coming, which have the capability to sequence multiple targets and samples simultaneously and can take advantage of a large number of Y-SNPs for forensic investigations on a detailed level. An early proof-of-principle study showed that 530 Y-SNPs could be genotyped simultaneously in a single sequencing run [25]. This Y-SNP NGS panel covered branches of the entire phylogenetic tree (Y-DNA Haplogroup Tree 2013) and could be applied to comprehensive paternal lineage classification. Subsequently, Ralf et al. presented a vastly improved Y-SNP NGS panel covering 859 Y-SNPs and 640 corresponding paternal lineages [34]. To promote the application of Y-SNPs for forensic investigations in China, multiple Y-SNP NGS panels aiming at the paternal lineage classification of Chinese populations were successively developed [27, 31, 35, 36]. However, Chinese populations possessed complex cultural, geographical, ethnic and genetic diversity patterns, these dedicated Y-SNP genotyping tools could not cover the dominant paternal lineages of Chinese populations and meet the requirement of high-resolution paternal lineage classification. Whole-genome sequencing-based genetic studies have illuminated Chinese population structures strongly correlated with the geographical regions or language families [37, 38]. Ancient DNA of East Eurasians further identified the westward steppe pastoralist ancestry, north-to-south di-directional population movements along the Yangtze River and Yellow River basins, peopling of the Tibetan Plateau and extensive interaction with ancient Siberians [39-43]. These ancient population events further complicated the patterns of the Y-chromosome lineages in China. To explore the fine-scale paternal genetic structure and illuminate the patterns of genetic diversity of Chinese populations, we developed one high-resolution revised Y-SNP panel with high coverage of geography and ethnicity specificity and the high resolution of terminal haplogroup dissection. We genotyped 639 Y-SNPs in 1033 unrelated individuals from 33 Han, Mongolian, Hui, Gelao, Li, and Manchu populations (**Figure 1A**). We comprehensively characterized the genetic diversity and population genetic features based on the sequence variations. We constructed one comprehensive revised forensic phylogenetic tree to present the patterns of Chinese Y-chromosome diversity and illuminate the high performance of our developed panel for forensic applications.

**Figure 1.**
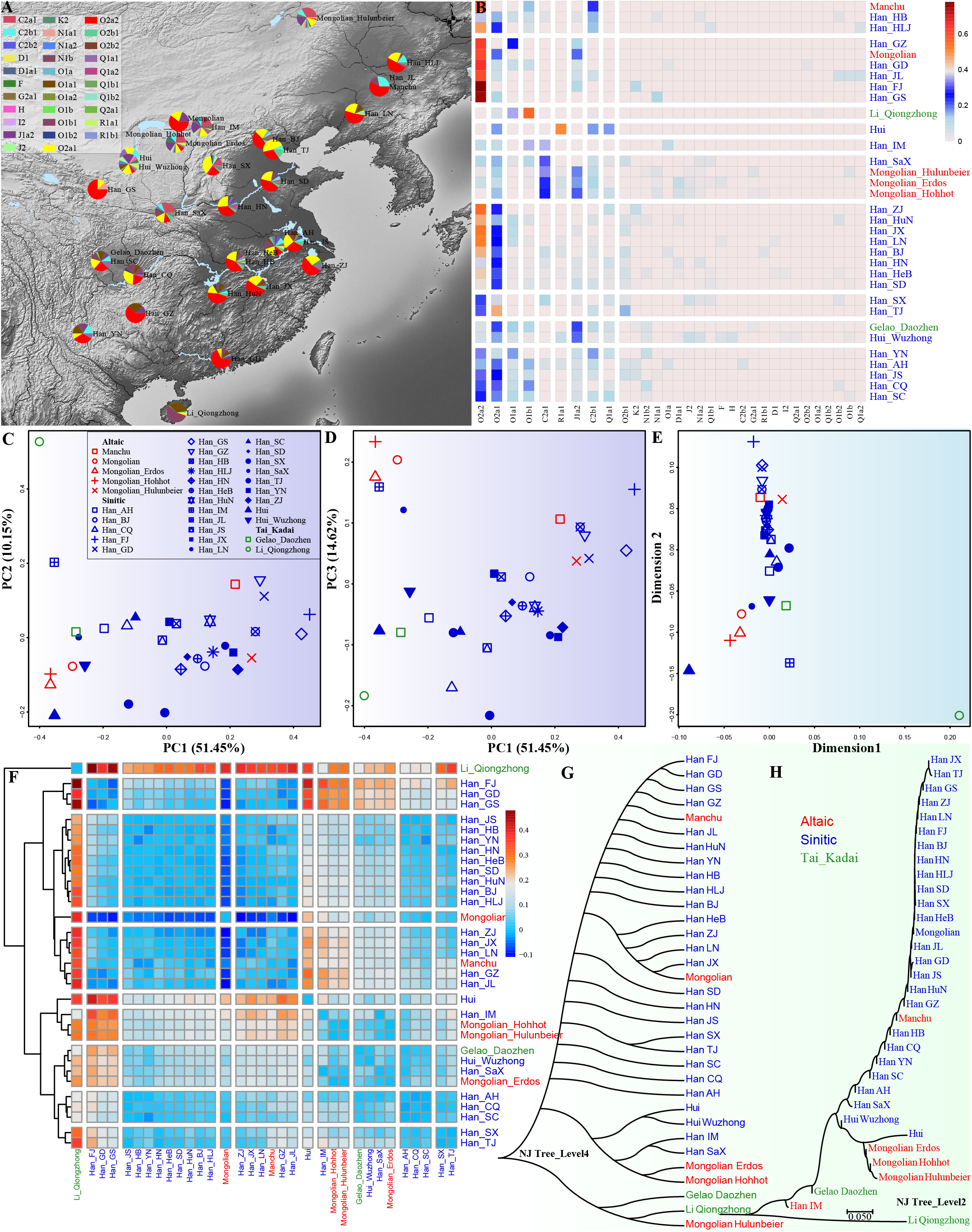
The geographical positions of 33 studied ethnolinguistically diverse populations and genetic features inferred from the haplogroup frequency spectrum. (**A**) Geographical locations and the haplogroup composition of 32 predefined 4-level haplogroups. All used haplogroups were manually cut at the fourth level to achieve a statistical possibility. (**B**) The heatmap showed the clustering patterns of 32 cut-haplogroups from 33 populations. (**C∼D**) Principal component analysis (PCA) showed the genetic similarities and differences among 33 populations based on the top three components extracted from the haplogroup frequency spectrum (HFS). (**E**) Multidimensional scaling plots showed the genetic clustering patterns based on the pairwise Fst matrix. (**F**) Heatmap showed the pairwise Fst calculated based on the HFS and the population clustered patterns (**G∼H**) Neighboring-Joining phylogenetic tree reconstructed based on the genetic distances in the different levels of HFS.

## MATERIALS AND METHODS

### Sample collection and DNA extraction

We collected peripheral blood samples from 1033 unrelated Han, Tibetan, Hui, Gelao, Manchu and Li individuals from 33 geographically different regions (**Figure 1A**). Each donor provided written informed consent. The medical ethics boards at Sichuan University and North Sichuan Medical College have approved our study protocol. Our experiments followed the recommendations and regulations of our institute and national guidelines of standards of the Declaration of Helsinki [44]. PureLink Genomic DNA Mini Kit (Thermo Fisher Scientific, Waltham, USA) was used to extract the genomic DNA. Based on the official manufacturer’s guidelines, Quantifiler Human DNA Quantification Kit (Thermo Fisher Scientific) and 7500 Real-time PCR System (Thermo Fisher Scientific) were used to quantify the DNA quantity and finally reserved in a low-temperature environment.

### Marker composition and NGS-based panel development

We chose the final included markers based on the following five rules to present a full-scale Chinese Y-chromosome diversity and high resolution of each terminal lineage. **First**, all major clade lineages recorded in the International Society of Genetic Genealogy (ISOGG) Y-DNA Haplogroup Tree 2019– 2020 version 15.73 (https://isogg.org/tree/index.html) and Yfull databases focused on Chinese populations were included. **Second**, key mutations included in our previously developed panel and validated in the population genetic studies were included [27, 45]. **Third**, we determined the terminal mutations based on the population-scale haplogroup frequency spectrum (HFS) according to the revised phylogenetic tree in the whole-genome sequencing projects. Based on the public data from the expanded 1000 Genomes Project cohort [3], Human Genetic Diversity Project (HGDP) [2], Simons Genome Diversity Project [46], Estonian Biocentre Human Genome Diversity Panel (EGDP) [47] and 10000 Chinese Person Genomic Diversity Project (10K_CPGDP) and others, we have built one in-house Y-chromosome population database with detailed HFS distribution and estimated divergence times (will be published soon). We choose the terminal mutations with a frequency larger than 5%. **Fourth**, based on the representatives of the included individuals and haplogroup coverage, we estimated the divergence times of each branch based on the localization of the mutations in the revised phylogeny. We included the mutations with a divergence time older than 500 years. We included 1000 Y-SNPs for the final primer design. We finally developed the Y-SNP NGS panel based on the MGISEQ-2000RS (MGI Tech Co., Ltd., Shenzhen, Guangdong, China) sequencing platform, which has been formally validated based on the SWGDAM guidelines (paper in preparation).

### Genotyping and quality control

We sequenced 639 Y-SNPs in 334 samples from Mongolian populations in three geographically different regions (159 males), Hui people from Wuzhong (57 males), Gelao people from Daozhen (59 males) and Li people (59 males) from Hainan, using our developed SNPSeqTyper 639 Y-SNP panel on the MGISEQ-2000RS sequencing platform. In each sequencing run, positive control of 9947 and negative ddH_2_O control were used. To provide one comprehensive comparative database, we also genotyped 639 Y-SNPs in 711 Han, Mongolian and Manchu individuals using an Affymetrix array. We used the PLINK v1.90b6.26 64-bit (2 Apr 2022) to conduct quality control on the merged dataset based on the missing SNP rate and missing genotyping rate with two parameters (--geno: 0.1 and --mind: 0.1) [48]. We finally kept a dataset of 639 Y-SNPs in 1033 unrelated individuals from 33 populations in the following forensic effectiveness evaluation and evolutionary history reconstruction.

### Classifying NRY haplogroups from VCF

We manually classified the NRY haplogroups for sequencing data based on the predefined phylogenetic tree with the chosen mutation markers. And then, we merged the sequencing data and the chip-based data and used the python package of hGrpr2.py instrumented in HaploGrouper to classify the haplogroups [49]. All branches in the haplogroup tree (treeFileNEW_isogg2019.txt) and snpFile_b38_isogg2019.txt were used in the HaploGrouper-based haplogroup classification. We additionally used the chip version (--chip) in the LineageTracker to classify the NRY haplogroups based on the GRCH38 reference genome [50].

### Haplogroup frequency spectrum estimation and clustering analysis

We calculated the HFS at different levels of the terminal haplogroups. We estimated the geographical distribution via a pie chart in the map and a heatmap based on the HFS matrix. Followingly, we conducted principal component analysis (PCA) based on the HFS matrix. We used the top three components extracted from the total variations to cluster our studied populations. We also calculated the pairwise Fst genetic matrix based on the HFS and conducted the multidimensional scaling analysis (MDS) to explore the genetic affinity between included populations.

### Phylogeny analysis for NRY haplogroups

We used the LineageTracker to construct the phylogenetic topology of all individuals [50]. Tools for variant calling and manipulating VCFs and BCFs (Bcftools Version: 1.8) and PLINK were used to convert the vcf files to fasta files. We used BEAUti to convert fasta files into XML files and ran the BEAST analysis using BEAST2.0 [51]. Tracer v1.7.2 was used to evaluate the power of statistical parameters and TreeAnnotator was used to choose the best trees in the BEAST results. We finally used FigTree v1.4.4 to visualize and organize the phylogenetic tree.

### Network analysis for Y-SNP haplotype data

Python package of fasta_to_nexus_Main.py was used to generate the nexus files. We used popART to construct the Network relationship. Here, five Network models were used to build the phylogenetic relationship among different lineages, including Minimum spanning network, Median Joining Network, Integer NJ Net, Tight Span Walker and TCS Network [52]. AMOVA was conducted to explore the genetic similarities and differences between or within groups and populations. Nucleotide diversity was also estimated using the popART [52].

## RESULTS

### Genotyping and genetic diversity among Chinese people

We generated genotype data of 639 Y-SNPs from 322 unrelated individuals from six Chinse populations belonging to three different language families using our developed SNPSeqTyper 639 Y-SNP panel, including three Mongolic-speaking Mongolian populations in North China, and one Sinitic-speaking Hui in Northwest China, two Tai-Kadai-speaking Gelao in Southwest China and Li in southernmost China (**Figure 1A**). To provide comprehensive population reference data from geographically diverse Chinese populations, we genotyped approximately 30K Y-SNPs in 711 Han, Manchu, Mongolian and Hui people from 27 populations using a high-density array, which included our genotyped 639 Y-SNPs. The average sequencing depth was high enough for our quality control. Among the final dataset of 1033 individuals from 33 populations, we observed 256 definitive Y-chromosomal lineages with the haplogroup frequency ranging from 0.0010 to 0.0687. Eighty-seven haplotypes were observed only once among our dataset, mainly including lineages from Q, R, C, D and other rare O terminal lineages. If we defined a haplogroup frequency larger than 5% as the threshold of the common lineages, the threshold of 1% as low-frequency lineages and other non-singleton as the rare lineages, we observed one common lineage (O2a2b1a1a1a1a1a1a1-M6539, 6.87%, 71/1033), 18 low-frequency lineages (O2a1b1a1a1a1a1a1-F17, O2a2b1a1a1a1a1b1a1b-MF15397, O2a2b2a1b1-A16609, O1b1a1a1a1b2a1a1-F2517, O2a2b1a1a1a1a1a1-F155, O1a1a1b-Z23420, C2a1a2a2b-F12502, O2a1b1a1a1a1-F325, O2a1b1a2a1a1-F1894, O2a2b1a2a1a1a2a1-F273, O2a1b1a1a1a1a1-F110, C2a1a3a-F3796, C2a1a1b1a-F3830, O2a1b1a1a1a1e1a-Y16154, J2-PF4922, Q1a1a1-F1626, O1a1a2a1a-Z23266 and O2b1a1a1-Y173834, the observed number over 11) and 150 rare lineages. We also statistically estimated the distribution of upstream Y-chromosomal lineages. We randomly cut the fourth level of each included haplogroup and observed six common lineages, including C2b1-F2613, O1a1a-CTS4351, C2a1-F3914, O1b1-F2320, O2a1-CTS7638 and O2a2-P201, seven low-frequency lineages, 15 rare lineages and sixvsingletons. Finally, we calculated the nucleotide diversity (π) among 302 individuals from six populations and obtained an estimate of 0.033. The estimated segregating sites of these included SNPs were 256 (the number of sites). The number of sites of parsimony-informative sites was 193. Tajima’s D statistic was -1.85776 with a p-value of 0.9856.

### Haplogroup frequency spectrum among 33 Chinese populations suggested their differentiated paternal genetic structure

We explored similarities and differences in the HFS among 33 investigated populations at level 4 (**Figure 1A∼B**). O1a only existed in Han Chinese, and the common lineage of O1a1 (0.3559) and low-frequency lineage of O1a2 (0.0169) were observed in Tai-Kadai-speaking Li with the highest proportion. Interestingly, Li-dominant lineage O1a1 was commonly identified in southern Han Chinese, Gelao and Hui people. O1b1, with the highest frequency of 0.5593, was mainly observed in southern Han Chinese populations and also identified in northern Han and Mongolian people. Two of four O2 lineages (O2a1 and O2a2) were frequently observed in Han Chinese. O2a1 had the highest haplogroup frequency in Tianjin Han, but O2a2 had the highest frequency in southern Hans (Fujian, Guangdong and Guizhou). C2a1 widely existed in Mongolian and other northern Han Chinese, and C2b1 was frequently observed in Manchu, Hui and other Han people. Except for the lineages mentioned above, we also observed that lineages of Q1a1, J1a2, R1a1 and R1b1 contributed to the paternal genetic diversity of Chinese populations (**Table S1** and **Figure 1A**). Heatmap of the HFS showed aforementioned common lineages contributed to the significant components of our studied population’s gene pool. We also found that geographically or ethnolinguistically close people shared similar patterns of HFS.

We additionally explored the genetic similarities and differences among 33 Chinese populations using PCA based on the HFS on the fourth level. PC1, extracting 51.45% variance from total variations, separated the northern and southern Chinese populations and PC2, extracting 10.15% variance, separated Tai-Kadai-speaking Li from other Chinese people (**Figure 1C**). Clustering patterns based on the first and third components separated Mongolian populations from others. Interestingly, Han Chinese populations from Inner Mongolia and Liaoning clustered closely with Mongolian people, suggesting their admixture and extensive interaction status (**Figure 1D**). Mongolian and Manchu people from the metropolitan populations clustered with Han Chinese suggested that southern Altaic people mixed with Han Chinese and other indigenous populations in the historical periods. These observed patterns were consistent with the admixture patterns inferred in our recent genomic studies of Mongolian and Manchu people from Guizhou province [53]. We further validated the identified patterns based on the pairwise Fst matrix, in which Li separated from others and Mongolian and northern Han clustered together and separated from others. Indeed, the estimated Fst values showed that Li people had the most considerable genetic distances with other comparative populations (**Table S3**) and separated from other populations, and formed one isolated clade in the heatmap clustering pattern (**Figure 1F**). Other populations were distinguished into two groups, mainly from northern and southern China. Finally, to confirm the robustness of our reconstructed genetic affinity, we reconstructed two Neighboring-Joining trees based on the Fst matrixes at the fourth and second levels (**Figure 1G∼H**). We found that the phylogenetic relationship inferred from the upstream Y-chromosome lineages was more consistent with the cluster patterns observed in the PCA, MDS and heatmap.

### High-resolution Y-chromosomal lineages for Non-Han Chinese populations

Recent whole-genome sequencing studies of Chinese populations have identified the fine-scale paternal genetic structure of ethnolinguistically different Chinese people and unreported LISNPs [10, 22, 23, 54]. However, whole-genome sequencing for every forensic case sample is impossible. Thus, our developed high-resolution Y-SNP panel is the best choice for promoting forensic applications. We chose all essential lineages in Chinese populations with the divergence times before 500 years. To validate the lineage coverage of our panel, we first genotyped 322 unrelated samples from six populations (Mongolian, Hui, Gelao and Li, **Figure 2A**). the reconstructed revised phylogeny among six ethnic groups showed that paternal lineages fell into O2a2, O2a1, O1b1, C2a1, O1a1, C2b1, Q1a1, R1a1 and D1a1, respectively sampled from 61, 46, 43, 42, 32, 24, 11, 10 and 9 individuals (**Figure 2A**). Dominant sublineages (O1b1a1a1a1b2a1a1, 23; O1a1a1b, 12 and C2a1a3a, 11) were observed and restricted to Li and Mongolian populations. Phylogeny results suggested that the C2a/2b can be identified as the founding lineage and ethnicity-specific lineage informative Y-SNP markers of Mongolians for further population genetics and forensic pedigree search as well as biogeographic ancestry inference. Similarly, the identified common sublineages of O1a/1b can be used as the Li-specific founding lineage for subsequent application. We also found some rare lineages originated from Siberia or western Eurasia and participated in the formation of Mongolian and Hui people in North China, suggesting the extensive population admixture along the population migration between North China, Siberia and Central Asia along the silk road or ancient Trans-Eurasian cultural and population communication. Based on the phylogenetic topology, we found that these founding lineages experienced population expansion and the admixture-introduced lineages remain a limited population size in Chinese populations.

**Figure 2.**
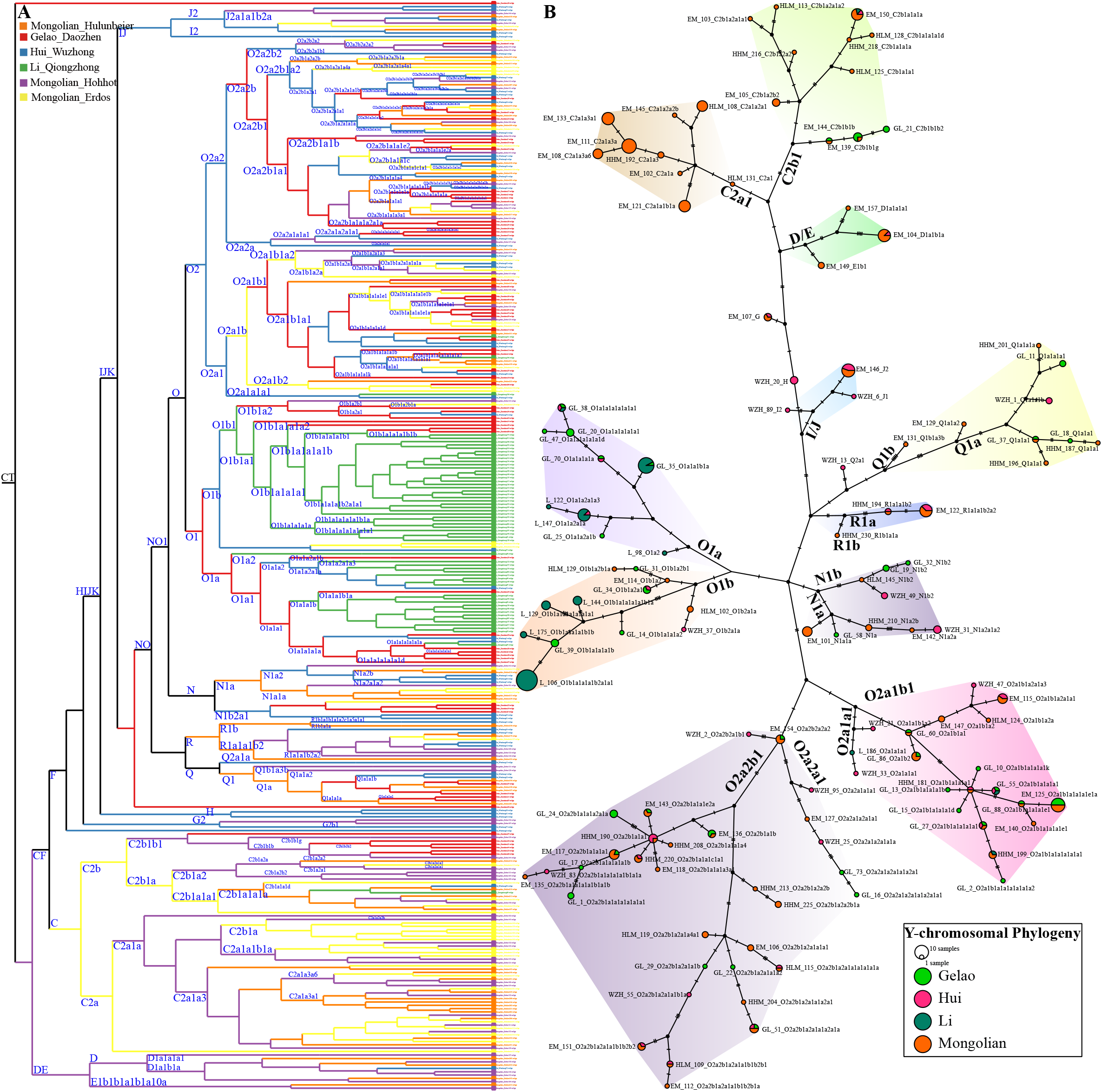
Phylogeny reconstruction among 322 Chinese populations from Mongolian, Gelao, Hui and Li people. (**A**) Phylogenetic relationships were reconstructed based on the Y-LineageTracker. Different colors denoted the diverse populations. (**B**) The Network reconstructed via the popART showed the shared haplotypes and mutations among other terminal haplogroups. The different colors showed different populations. Different color backgrounds denoted the highlighted Y-chromosomal lineages among Chinese populations.

To directly explore the genetic connection among different Y-chromosomal lineages and ethnically different populations, we constructed the Network relationship among 322 Chinese males. Consistent with the identified common, rare, or low-frequency lineages in the phylogenetic topology, we observed eleven different lineages contributed to the genetic diversity of these studied populations. C2a1 lineage was unique in Mongolian, and C2b1 was dominant in Mongolian and presented some sublineages in Gelao. O1a/b was dominant in southern Chinese populations (Gelao and Li). O2a2a1/b1 and O2a1a1, O2a1a1/b1 were contributed to Mongolian, Hui and Gelao. We could also identify some star-like expansion of some dominant lineages, such as C2a2b1a1a1 in Mongolian populations.

### Extensive population expansion and admixture within and between Han Chinese populations and other minorities

Han Chinese is the largest ethnic group in the world and has a significant influence on the formation of the gene pool of Chinese populations [37, 38, 55-57]. To further explore phylogenetic topology among Han Chinese and illuminate the genetic relationship between Han Chinese and aforesaid minorities, we collected and genotyped 711 individuals via Affymetrix array, which included 690 Han Chinese from 25 geographically different populations. We reconstructed a more representative phylogeny using 1033 sequences and confirmed the Mongolian and Li-specific founding lineages (**Figure 3**). We found that these two founding lineages participated in the formation of Han Chinese populations. Among Han Chinese populations, we also identified the western Eurasian-introduced R1a/1b and Siberian-mediated Q1. We identified Han Chinese dominant Y-chromosome lineages of O2a2, O2a1, O1b1, O1a1, C2b1, C2a1, O2b1and Q1a1, respectively sampled from 303, 147, 38, 35, 33, 26, 26 and 18 individuals. Some sublineages were also underwent recent population expansion (O2a2b1a1a1a1a1a1a1, 69; O2a1b1a1a1a1a1a1, 38; O2a2b1a1a1a1a1b1a1b, 5; O2a2b1a1a1a1a1a1, 22; O2a2b2a1b1, 22). The observed mosaic patterns of identified paternal lineages showed extensive gene flow among Han Chinese populations and minority groups, and the gene flow influence was bi-directional. Mongolian and Li dominant founding lineages were observed in Han Chinese individuals, suggesting that ancient Baiyue ancestors and Eurasian pastoralist people participated in the formation of Han Chinese. Han Chinese dominant lineages were also identified in Mongolian, Hui, Li and Gelao people, which supported population interaction from the paternal perspective.

**Figure 3.**
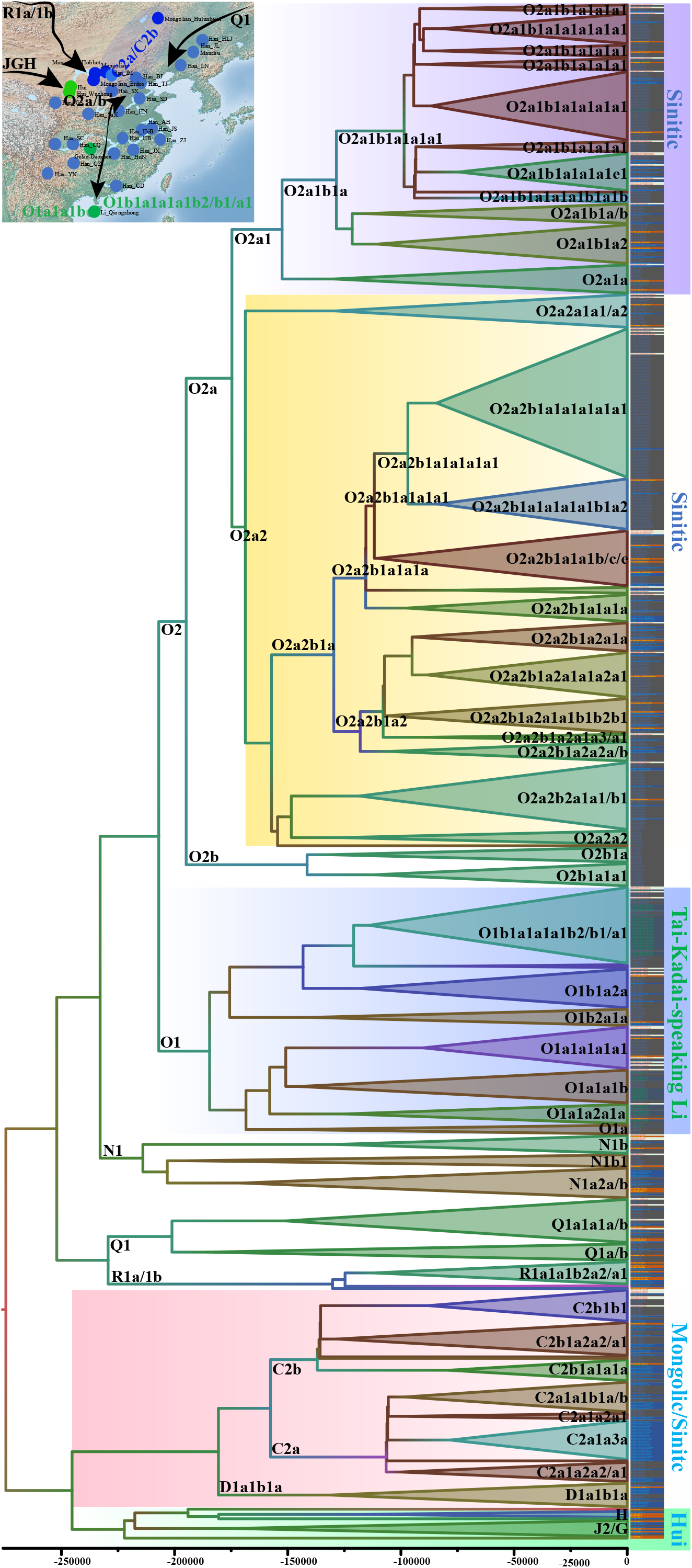
Phylogenetic relationships based on the sequence diversity among 1033 Chinese males from 33 populations. Different color in a circle showed their different composition of population origins. The different color backgrounds showed different founding lineages. The haplogroup followed by the sample ID was classified using the HaploGrouper.

The reconstructed merged Network confirmed the complex admixture and expansion events among Han Chinese and other non-Han Chinese populations (**Figure 4**). The Mongolian and Li-specific lineages (C2a/b and O1a/b) were separated from the Han Chinese-related lineages. We could also find that Li and Mongolian dominant lineages influenced the Han Chinese gene pool as the population composition of one target haplogroup. The distribution of samples belonging to O2a1/a2 denoted that their primary ancestry was derived from Han Chinese populations and minor ancestry from Li. But the obtained ancestral lineages from Han Chinese in Gelao, Mongolian, and Hui people were more evident than that in Li people. O2a2b1a1a1a1 and O2a1b1a1a1a1e1 were two important lineages that experienced population expansion.

**Figure 4.**
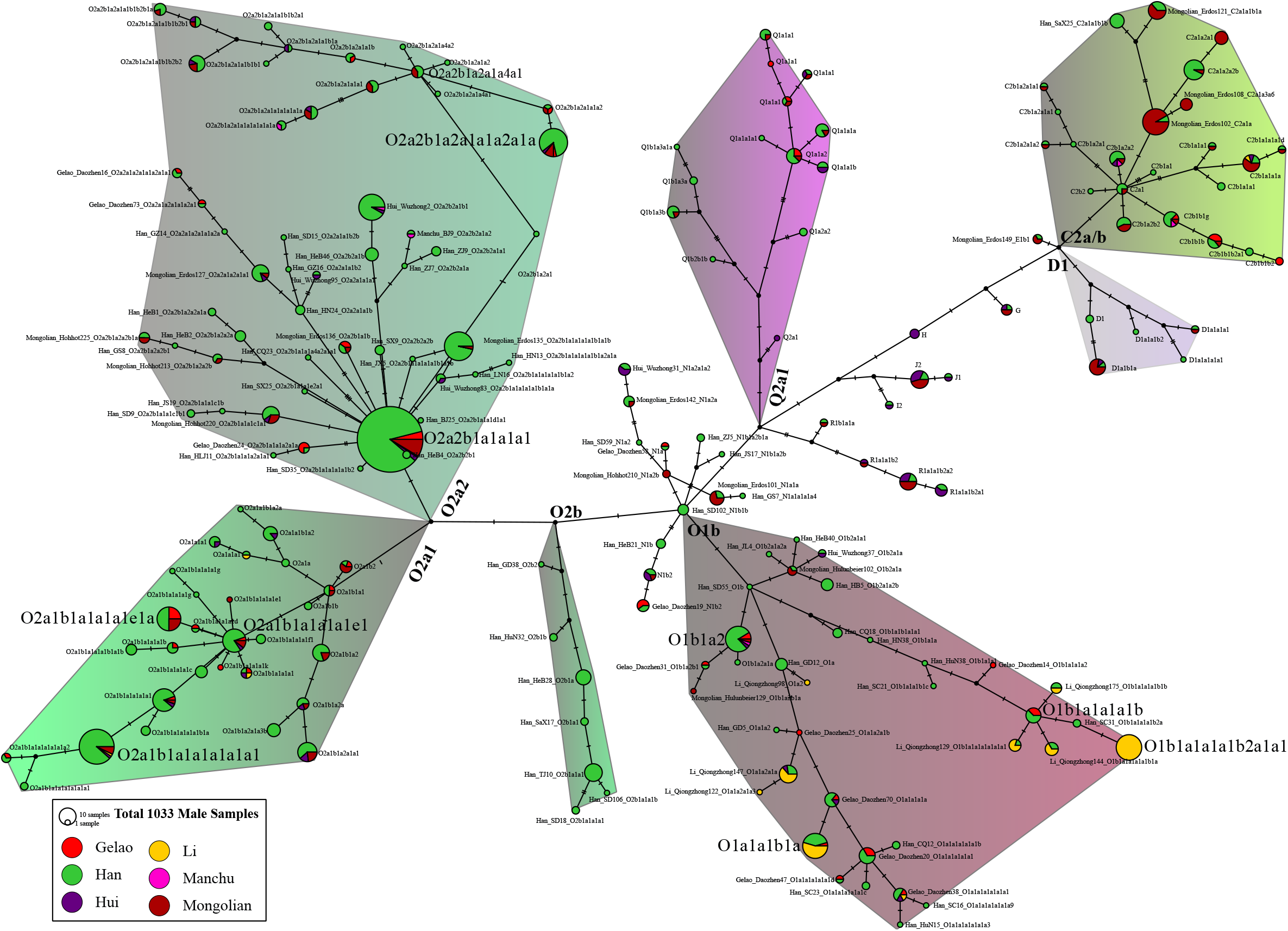
The reconstructed phylogeny of Chinese populations. The map showed the primary distribution of our investigated Y-chromosomal lineages. The arrow showed the possible migration direction. Circles in the map were coded based on their language families. The line color and triangle denoted the posterior. The triangle length showed the population size and the height showed the relative divergence time. Population ID was categorized based on their ethnicity belongs.

## DISCUSSION

### The full landscape of Y-chromosomal diversity reveals complex population migration and admixture tracts

Non-crossover regions of the human Y-chromosome harbor the feature of male-specific inheritance and can record most male behavior, phenotype and human demographic details [4, 7]. To explore the patterns of Y-chromosomal diversity, we reported the genotypes of 1033 Y chromosomes randomly sampled from 33 Chinese populations belonging to five ethnic groups (Mongolian, Manchu, Hui, Gelao, Li and Han), which were genotyped using our newly-developed SNPSeqTyper 639 Y-SNP panel and high-density Affymetrix array. We have conducted a comprehensive population evolutionary analysis and population comparison tests within and between Chinese populations belonging to different geographical regions or language families. Population genetic survey suggested that our panel captured the richest Y-chromosomal genetic diversity to date in all forensic Y-SNP genotyping tools focused on the Chinese populations [27, 32, 45, 58-62]. Phylogeny constructed among Non-Han Chinese (Mongolic, Hui and Tai-Kadai) and all included subjects consistently demonstrated that strong geography or ethnicity-related Y-chromosomal features indicated the underlying complex population evolutionary history and potential for forensic pedigree search and biogeographical ancestry inference.

HFS analysis among our studied populations or regional northern Mongolian and southern island Li people has revealed their common founding lineages of C2a/2b and O1a/1b (**Figure 1A∼B**). Geographical distribution further confirmed these dominant lineages could be used as forensic markers for genetic localization. C2a1-F3914 can be used as a Mongolian predominant founding lineage, which was observed in 42 Mongolian individuals and 26 Han Chinese individuals, mainly from Shaanxi, Shanxi and Inner Mongolia. Another Mongolian predominant lineage C2b1-F2613 was observed in 16 Mongolians (16/159), two Manchu, one Li, two Hui, 32 Han (32/693) and six Gelao individuals. Upstream C2b-F1067 (previously classified as C2c) was reported first in northeastern Asia and associated with the origin and expansion of Mongolic-speaking populations. Subclades of C2b1a1a1a-M407 (C2c1a1a1 in the previous version) appeared in ten individuals (two Hans, one Hui, one Li and four Mongolians), and C2b1b1b-F5477 and C2b1a2b2-FGC45548 were also respectively observed seven and six times. Huang et al. presented one revised phylogenetic tree and distribution map focused on all available C2b1a1a1a-M407 samples and found that C2b1a1a1a-M407 has a frequency of over 50% in northeastern Asian population [63]. Thus, C2b1-F2613 and C2b1a1a1a-M407 can be used to trace the population origin and migration of Kalmyks, Mongolians, Buryats and other genetically close northeastern Asians.

Network and phylogeny constructions further showed that the lineage of O1a-M119 was the common paternal lineage in southern Chinese populations (**Figure 2**). Sublineages of O1b1a1a1a1b2a1a1-F2517 (23), O1b1a2a1-F1759 (10), O1a1a1b-Z23420 (15) and O1a1a2a1a-Z23266 (11) had undergone population expansion recently. All F2517 lineages were observed in Li people, which was consistent with a recent whole-genome sequencing study [64]. Chen et al. found O1b1a1a was the dominant lineage in southern East Asians, which diverged from others 10998 years ago, and the F2517 sublineages were further divided into O1b1a1a1a1b2a1a1a and O1b1a1a1a1b2a1a1b clades at 2828 years ago [64]. Sun et al. also found most sublineages of M119, including our identified F1759, Z23420 and Z23266, contributed to the ancient gene pool of modern Tai-Kadai, Austronesian and southern Han Chinese [10]. We could also identify the shared paternal lineages among Li, southern Han and Gelao people in the distribution of M119 mutations. The unique paternal genetic structure of the Hainan Li people was also consistent with our previous findings of the fine-scale genetic structure [65].

Our survey has identified 195 samples of O2a1-CTS7638 and 372 samples of O2a2-P201 in Han Chinese populations. Sublineages of O2a2b1a1a1a1a1a1a1-M6539 (71), O2a1b1a1a1a1a1a1-F17 (43), O2a2b1a1a1a1a1b1a1b-MF15397 (26), O2a2b2a1b1-A16609 (24) and O2a2b1a1a1a1a1a1-F155 (23) have experienced population expansion events recently, which was consistent with population expansion among Han Chinese populations inferred from the whole-genome sequencing project. Admixture-introduced rare lineage Q1a1-F746/F790 was observed in 30 samples from Mongolian, Han, and Gelao people, and steppe pastoralist-related R1a1a1b2-Z93 was observed in 17 Mongolian and Hui people. Our results provided genetic evidence for extensive admixture between northern East Asians and surrounding populations. Similar patterns were also illuminated in the paternal genetic history reconstruction by Wang et al., who concluded that multiple ancestral sources contributed to the formation of paternal gene pool of Mongolian people [32]. Besides, He et al. found that Mongolic-speaking populations have strong population stratifications, in which the northern one was influenced by Siberian ancestry, the western one was influenced by western Eurasian and the southern one was influenced by Han Chinese expansion [37]. Recent ancient DNA also found that western Eurasian steppe ancestry has influenced the genetic makeup of northern East Asians. For western Eurasian ancestry identified in Hui people, our paternal results were also consistent with the admixture patterns via the genome-wide SNP data. Complex demographical models suggested that geographically diverse Chinese Hui people harbored complex and different admixture processes and possessed approximately 10% ancestry related to the ancestor from Central Asians [66-68].

We also identified paternal genetic structure among Chinese populations in the clustering patterns via HFS, PCA and MDS. These population stratifications were in accordance with the language or geographical categories. Island Li people shared their specific paternal genetic structure and clustered far from other Chinese populations. Similarly, Mongolian people clustered together and have a relatively close genetic relationship with northern East Asians. We also should note that the Y-chromosome-based population structure in China is rougher than that inferred from the genome-wide SNPs. Our recent genetic studies have identified five population substructures correlated with languages and geography in China. Mongolic and Tungusic people in Northeastern China harbored the highest Ulchi or ancient Boisman/DevilsCave-related ancestry [37, 41]. Tibeto-Burman groups from Tibetan Plateau had the highest proportion of ancestry related to core Tibet Tibetan and Nepal Chokhopani, Mebrak, and Samdzong people[69, 70]. The primary ancestral component of Hmong-Mien people from southwestern China was maximized in ancestry related to Miao and Yao people [71], and Austronesian-speaking people from Taiwan Island have more Ami/Atayal or ancient Hanben-related ancestry [41]. Han Chinese ancestry localized between the four ancestries mentioned above and showed a northern-to-southern genetic cline. The recent large-scale genetic structure also identified fine-scale population structure among geographically diverse Han Chinese populations [55, 56, 72]. These fine-scale genetic backgrounds could promote better study design for large medical clinical cohorts and forensic genetic localization of crime cases.

### SNPSeqTyper 639 Y-SNP panel can be used as a powerful forensic tool for Chinese forensic pedigree search and biogeographic ancestry inference

The forensic community has noticed that whole-genome sequencing in a forensic case needs to overcome specific infrastructure of the specialist, platform and genomic statistical methods, as well as the experiment method focused on the forensic case samples. Besides, the cost of one sample is another important obstacle to the wide application of whole-genome sequencing technology in forensics. Evolutionary genetic scientists have conducted many vital projects to explore the complete anthropologically-informed phylogeny [7, 19, 20]. Our work has identified most paternal founding lineages in Chinese populations and comprehensively characterized their geographical and ethnic distribution. Our panel harbored the high coverage of genetic variations of terminal lineages and complete coverage of reference data from main populations or ethnic groups in China.

Forensic pedigree search can help trace possible crime suspects based on the shared Y-chromosome mutations. Lineages informative Y-SNPs were usually used together with Y-STR markers. Many prior works provided relatively high-resolution forensic phylogenic trees and presented the corresponding scientific examination and analysis strategies. They promoted the advances of forensic Y-chromosome applications in pedigree search and biogeographic ancestry inference based on the customized SNaPshot and NGS technologies [27, 32, 45, 58-62]. Song et al. explored the paternal genetic structure of Hainan Li using their developed panel containing 141 Y-SNPs. They found haplogroup O1b1a1a1a1a1b-CTS5854 can be used as one ethnicity-specific lineage in population and forensic genetics [61]. Song et al. further updated their panel, including 233 Y-SNPs used for Chinese Qiang people, and found that O2a2b1a1-M117, O2a2b1a1a1-F42 and O2a1b1a1a1a-F11 were the founding lineages in Qiang people [62]. Wang et al. also investigated the paternal genetic structure of Zhuang people using this panel and identified the O2-dominant lineages in Tai-Kadai people [59]. Xie et al. developed one panel focused on Hui people, which included 157 Y-SNP, and identified the population substructure of Hui people [58]. Wang et al. focused on the genetic diversity of Mongolian people (N1b-F2930, N1a1a1a1a3-B197, Q-M242, and O2a2b1a1a1a4a-CTS4658) and developed one Mongolian-specific panel included 215 Y-SNPs [32]. The panels mentioned above consisted of several customized SNaPshot systems, which limited the quakily used in forensic cases. Wang et al. developed one 165-plex Y-SNP panel based on an Ion S5 XL system. They comprehensively conducted the sequencing performance and concordance, reliability, sensitivity, and stability studies based on the ISFG guidelines [27]. Liu et al. recently updated this system by increasing the final Y-SNP number to 256 [45]. Significantly, Tao et al. developed a customized SifaMPS 381 Y-SNP panel that included 381 Y-SNPs focused on Chinese populations and investigated the basic structure and subbranches of Chinese major haplogroup branches [60]. Our panel included two significant features: the first one is lineages specific to or common in most Chinese populations were included (O, D, C, R, and Q et al.), and the other important one is that we retained a higher resolution of the terminal Y-chromosome lineages, which can complete the shortcoming of previously developed panels limited to some common lineages or only focused on specific populations. The newly-developed panel overcame the limitation of the lineage’s representatives, terminal lineage resolution and sequencing platforms, which can provide the best practice tool in forensic applications. The identified paternal population structure in China can provide more clues for biogeographical ancestry inference, which can be used as a complementary tool for forensic ancestry prediction based on the autosome-based ancestry informative SNP panel [66].

## CONCLUSION

The complete landscape of human Y-chromosome variations and the gradually updated Y-chromosome phylogenetic tree with more population-specific LISNPs formed the fundamental for forensic application and evolutionary study. To overcome the shortness of the whole Y-chromosome sequencing in forensic science at the initial stage of the genome sequencing era, we developed one highest-resolution SNPSeqTyper 639 Y-SNP panel that included 639 LISNPs defined 574 terminal Y-chromosomal lineages. We generated the population data from 1033 individuals from 33 populations, including Han, Hui, Mongolian, Li and Gelao and investigated the forensic features, HFS and evolutionary processes via multiple statistical models. We identified 257 terminal Y-chromosomal lineages with several common founding lineages of O2a2b1a1a1a1a1a1a1-M6539, O2a1b1a1a1a1a1a1-F17, O2a2b1a1a1a1a1b1a1b-MF15397, O2a2b2a1b1-A16609, O1b1a1a1a1b2a1a1-F2517 and O2a2b1a1a1a1a1a1-F155. Patterns of HFS and corresponding geographical distribution illuminated that some Siberian or western Eurasian-originated paternal lineages contributed to the formation of the paternal gene pool of Mongolian, Hui and northern Han Chinese populations. Network and our reconstructed forensic phylogenic topology further illuminated the complex population divergence and expansion of different paternal lineages, which also found some ancestral lineages shared by geographically or linguistically different Chinese populations. PCA and MDS clustering patterns showed that the paternal genetic structure was correlated with the geographical and linguistic categories, which provided the basic genetic background for forensic paternal biogeographical ancestry inference. Our reconstructed revised phylogeny and comprehensive population genetic investigation based on this Y-SNP panel can provide the highest resolution of the terminal lineage and genetic diversity, which provides one panel with high marker coverage and lineage representative. Ethnolinguistically diverse Chinese populations had the highest genetic diversity. Thus, anthropologically-informed Y-chromosome whole-genome sequencing will promote the further development of higher-resolution Y-SNP NGS panels and corresponding population-specific dataset construction. We also emphasized that the large-scale population cohorts, such as 10K_CPGDP and 100KGSRD^WCH^ (100K genome sequencing of rare disease), can provide more unreported LISNPs for forensic application of Y-chromosome.

## ACKNOWLEDGMENTS

This work is funded by the National Natural Science Foundation of China (82202078) and the Open project of Key Laboratory of Forensic Genetics of Ministry of Public Security (2022FGKFKT05).

## DATA AVAILABILITY

We submitted allele frequency data in the public database <u>(Available when it is publised online)</u>. Following this project’s regulations, informed consent, and the Human Genetic Resources Administration of China (HGRAC) regulations, we can share the obtained genotype data via personal communication with the corresponding authors. We make the data available upon request by asking the person requesting the data to agree in writing to the following restrictions: 1, the data can be only used for studying population history; 2, the data cannot be used for commercial purposes; 3, the data cannot be used to identify the sample donors; 4, the data cannot be used for studying natural/cultural selections, medical or other related studies.

## REFERENCES

[1] M.A. Almarri, A. Bergström, J. Prado-Martinez, F. Yang, B. Fu, A.S. Dunham, Y. Chen, M.E. Hurles, C. Tyler-Smith, Y. Xue, Population Structure, Stratification, and Introgression of Human Structural Variation, Cell 182(1) (2020) 189-199.e15.

[2] A. Bergstrom, S.A. McCarthy, R. Hui, M.A. Almarri, Q. Ayub, P. Danecek, Y. Chen, S. Felkel, P. Hallast, J. Kamm, H. Blanche, J.F. Deleuze, H. Cann, S. Mallick, D. Reich, M.S. Sandhu, P. Skoglund, A. Scally, Y. Xue, R. Durbin, C. Tyler-Smith, Insights into human genetic variation and population history from 929 diverse genomes, Science 367(6484) (2020).

[3] M. Byrska-Bishop, U.S. Evani, X. Zhao, A.O. Basile, H.J. Abel, A.A. Regier, A. Corvelo, W.E. Clarke, R. Musunuri, K. Nagulapalli, S. Fairley, A. Runnels, L. Winterkorn, E. Lowy, C. Human Genome Structural Variation, F. Paul, S. Germer, H. Brand, I.M. Hall, M.E. Talkowski, G. Narzisi, M.C. Zody, High-coverage whole-genome sequencing of the expanded 1000 Genomes Project cohort including 602 trios, Cell 185(18) (2022) 3426–3440 e19.

[4] M.A. Jobling, C. Tyler-Smith, Human Y-chromosome variation in the genome-sequencing era, Nat Rev Genet 18(8) (2017) 485–497.

[5] M. Kayser, Forensic use of Y-chromosome DNA: a general overview, Hum Genet 136(5) (2017) 621–635.

[6] H. Yao, S. Wen, X. Tong, B. Zhou, P. Du, M. Shi, L. Jin, H. Li, Y chromosomal clue successfully facilitated the arrest of Baiyin serial killer, Science Bulletin 61(22) (2016) 1715–1717.

[7] G.D. Poznik, Y. Xue, F.L. Mendez, T.F. Willems, A. Massaia, M.A. Wilson Sayres, Q. Ayub, S.A. McCarthy, A. Narechania, S. Kashin, Y. Chen, R. Banerjee, J.L. Rodriguez-Flores, M. Cerezo, H. Shao, M. Gymrek, A. Malhotra, S. Louzada, R. Desalle, G.R. Ritchie, E. Cerveira, T.W. Fitzgerald, E. Garrison, A. Marcketta, D. Mittelman, M. Romanovitch, C. Zhang, X. Zheng-Bradley, G.R. Abecasis, S.A. McCarroll, P. Flicek, P.A. Underhill, L. Coin, D.R. Zerbino, F. Yang, C. Lee, L. Clarke, A. Auton, Y. Erlich, R.E. Handsaker, C. Genomes Project, C.D. Bustamante, C. Tyler-Smith, Punctuated bursts in human male demography inferred from 1,244 worldwide Y-chromosome sequences, Nat Genet 48(6) (2016) 593–9.

[8] M. van Oven, A. Van Geystelen, M. Kayser, R. Decorte, M.H. Larmuseau, Seeing the wood for the trees: a minimal reference phylogeny for the human Y chromosome, Hum Mutat 35(2) (2014) 187–91.

[9] P. Hallast, C. Batini, D. Zadik, P. Maisano Delser, J.H. Wetton, E. Arroyo-Pardo, G.L. Cavalleri, P. de Knijff, G. Destro Bisol, B.M. Dupuy, H.A. Eriksen, L.B. Jorde, T.E. King, M.H. Larmuseau, A. Lopez de Munain, A.M. Lopez-Parra, A. Loutradis, J. Milasin, A. Novelletto, H. Pamjav, A. Sajantila, W. Schempp, M. Sears, A. Tolun, C. Tyler-Smith, A. Van Geystelen, S. Watkins, B. Winney, M.A. Jobling, The Y-chromosome tree bursts into leaf: 13,000 high-confidence SNPs covering the majority of known clades, Mol Biol Evol 32(3) (2015) 661–73.

[10] J. Sun, Y.X. Li, P.C. Ma, S. Yan, H.Z. Cheng, Z.Q. Fan, X.H. Deng, K. Ru, C.C. Wang, G. Chen, L.H. Wei, Shared paternal ancestry of Han, Tai-Kadai-speaking, and Austronesian-speaking populations as revealed by the high resolution phylogeny of O1a-M119 and distribution of its sub-lineages within China, Am J Phys Anthropol 174(4) (2021) 686–700.

[11] W. Kutanan, J. Kampuansai, M. Srikummool, A. Brunelli, S. Ghirotto, L. Arias, E. Macholdt, A. Hubner, R. Schroder, M. Stoneking, Contrasting Paternal and Maternal Genetic Histories of Thai and Lao Populations, Mol Biol Evol 36(7) (2019) 1490–1506.

[12] T. Zerjal, Y. Xue, G. Bertorelle, R.S. Wells, W. Bao, S. Zhu, R. Qamar, Q. Ayub, A. Mohyuddin, S. Fu, P. Li, N. Yuldasheva, R. Ruzibakiev, J. Xu, Q. Shu, R. Du, H. Yang, M.E. Hurles, E. Robinson, T. Gerelsaikhan, B. Dashnyam, S.Q. Mehdi, C. Tyler-Smith, The genetic legacy of the Mongols, Am J Hum Genet 72(3) (2003) 717–21.

[13] Y. Xue, T. Zerjal, W. Bao, S. Zhu, S.K. Lim, Q. Shu, J. Xu, R. Du, S. Fu, P. Li, H. Yang, C. Tyler-Smith, Recent spread of a Y-chromosomal lineage in northern China and Mongolia, Am J Hum Genet 77(6) (2005) 1112–6.

[14] B. Wen, H. Li, D. Lu, X. Song, F. Zhang, Y. He, F. Li, Y. Gao, X. Mao, L. Zhang, J. Qian, J. Tan, J. Jin, W. Huang, R. Deka, B. Su, R. Chakraborty, L. Jin, Genetic evidence supports demic diffusion of Han culture, Nature 431(7006) (2004) 302–5.

[15] W. Wei, Q. Ayub, Y. Chen, S. McCarthy, Y. Hou, I. Carbone, Y. Xue, C. Tyler-Smith, A calibrated human Y-chromosomal phylogeny based on resequencing, Genome Res 23(2) (2013) 388–95.

[16] M. Karmin, L. Saag, M. Vicente, M.A. Wilson Sayres, M. Jarve, U.G. Talas, S. Rootsi, A.M. Ilumae, R. Magi, M. Mitt, L. Pagani, T. Puurand, Z. Faltyskova, F. Clemente, A. Cardona, E. Metspalu, H. Sahakyan, B. Yunusbayev, G. Hudjashov, M. DeGiorgio, E.L. Loogvali, C. Eichstaedt, M. Eelmets, G. Chaubey, K. Tambets, S. Litvinov, M. Mormina, Y. Xue, Q. Ayub, G. Zoraqi, T.S. Korneliussen, F. Akhatova, J. Lachance, S. Tishkoff, K. Momynaliev, F.X. Ricaut, P. Kusuma, H. Razafindrazaka, D. Pierron, M.P. Cox, G.N. Sultana, R. Willerslev, C. Muller, M. Westaway, D. Lambert, V. Skaro, L. Kovacevic, S. Turdikulova, D. Dalimova, R. Khusainova, N. Trofimova, V. Akhmetova, I. Khidiyatova, D.V. Lichman, J. Isakova, E. Pocheshkhova, Z. Sabitov, N.A. Barashkov, P. Nymadawa, E. Mihailov, J.W. Seng, I. Evseeva, A.B. Migliano, S. Abdullah, G. Andriadze, D. Primorac, L. Atramentova, O. Utevska, L. Yepiskoposyan, D. Marjanovic, A. Kushniarevich, D.M. Behar, C. Gilissen, L. Vissers, J.A. Veltman, E. Balanovska, M. Derenko, B. Malyarchuk, A. Metspalu, S. Fedorova, A. Eriksson, A. Manica, F.L. Mendez, T.M. Karafet, K.R. Veeramah, N. Bradman, M.F. Hammer, L.P. Osipova, O. Balanovsky, E.K. Khusnutdinova, K. Johnsen, M. Remm, M.G. Thomas, C. Tyler-Smith, P.A. Underhill, E. Willerslev, R. Nielsen, M. Metspalu, R. Villems, T. Kivisild, A recent bottleneck of Y chromosome diversity coincides with a global change in culture, Genome Res 25(4) (2015) 459–66.

[17] P. Francalacci, L. Morelli, A. Angius, R. Berutti, F. Reinier, R. Atzeni, R. Pilu, F. Busonero, A. Maschio, I. Zara, D. Sanna, A. Useli, M.F. Urru, M. Marcelli, R. Cusano, M. Oppo, M. Zoledziewska, M. Pitzalis, F. Deidda, E. Porcu, F. Poddie, H.M. Kang, R. Lyons, B. Tarrier, J.B. Gresham, B. Li, S. Tofanelli, S. Alonso, M. Dei, S. Lai, A. Mulas, M.B. Whalen, S. Uzzau, C. Jones, D. Schlessinger, G.R. Abecasis, S. Sanna, C. Sidore, F. Cucca, Low-pass DNA sequencing of 1200 Sardinians reconstructs European Y-chromosome phylogeny, Science 341(6145) (2013) 565–9.

[18] A. Bergstrom, N. Nagle, Y. Chen, S. McCarthy, M.O. Pollard, Q. Ayub, S. Wilcox, L. Wilcox, R.A. van Oorschot, P. McAllister, L. Williams, Y. Xue, R.J. Mitchell, C. Tyler-Smith, Deep Roots for Aboriginal Australian Y Chromosomes, Curr Biol 26(6) (2016) 809–13.

[19] T. Pinotti, A. Bergstrom, M. Geppert, M. Bawn, D. Ohasi, W. Shi, D.R. Lacerda, A. Solli, J. Norstedt, K. Reed, K. Dawtry, F. Gonzalez-Andrade, Y.M.C. Paz, S. Revollo, C. Cuellar, M.S. Jota, J.E. Santos, Jr., Q. Ayub, T. Kivisild, J.R. Sandoval, R. Fujita, Y. Xue, L. Roewer, F.R. Santos, C. Tyler-Smith, Y Chromosome Sequences Reveal a Short Beringian Standstill, Rapid Expansion, and early Population structure of Native American Founders, Curr Biol 29(1) (2019) 149–157 e3.

[20] M. Karmin, R. Flores, L. Saag, G. Hudjashov, N. Brucato, C. Crenna-Darusallam, M. Larena, P.L. Endicott, M. Jakobsson, J.S. Lansing, H. Sudoyo, M. Leavesley, M. Metspalu, F.X. Ricaut, M.P. Cox, Episodes of Diversification and Isolation in Island Southeast Asian and Near Oceanian Male Lineages, Mol Biol Evol 39(3) (2022) msac045.

[21] L.X. Wang, Y. Lu, C. Zhang, L.H. Wei, S. Yan, Y.Z. Huang, C.C. Wang, S. Mallick, S.Q. Wen, L. Jin, S.H. Xu, H. Li, Reconstruction of Y-chromosome phylogeny reveals two neolithic expansions of Tibeto-Burman populations, Mol Genet Genomics 293(5) (2018) 1293–1300.

[22] N. Sun, P.C. Ma, S. Yan, S.Q. Wen, C. Sun, P.X. Du, H.Z. Cheng, X.H. Deng, C.C. Wang, L.H. Wei, Phylogeography of Y-chromosome haplogroup Q1a1a-M120, a paternal lineage connecting populations in Siberia and East Asia, Ann Hum Biol 46(3) (2019) 261–266.

[23] B.L. Liu, P.C. Ma, C.Z. Wang, S. Yan, H.B. Yao, Y.L. Li, Y.M. Xie, S.L. Meng, J. Sun, Y.H. Cai, S. Sarengaowa, H. Li, H.Z. Cheng, L.H. Wei, Paternal origin of Tungusic-speaking populations: Insights from the updated phylogenetic tree of Y-chromosome haplogroup C2a-M86, American journal of human biology : the official journal of the Human Biology Council 33(2) (2021) e23462.

[24] M.A. Jobling, C. Tyler-Smith, The human Y chromosome: an evolutionary marker comes of age, Nat Rev Genet 4(8) (2003) 598–612.

[25] A. Ralf, M. van Oven, K. Zhong, M. Kayser, Simultaneous analysis of hundreds of Y-chromosomal SNPs for high-resolution paternal lineage classification using targeted semiconductor sequencing, Hum Mutat 36(1) (2015) 151–9.

[26] M. Xie, F. Song, J. Li, M. Lang, H. Luo, Z. Wang, J. Wu, C. Li, C. Tian, W. Wang, H. Ma, Z. Song, Y. Fan, Y. Hou, Genetic substructure and forensic characteristics of Chinese Hui populations using 157 Y-SNPs and 27 Y-STRs, Forensic Sci Int Genet 41 (2019) 11–18.

[27] M. Wang, Z. Wang, G. He, J. Liu, S. Wang, X. Qian, M. Lang, J. Li, M. Xie, C. Li, Y. Hou, Developmental validation of a custom panel including 165 Y-SNPs for Chinese Y-chromosomal haplogroups dissection using the ion S5 XL system, Forensic Sci Int Genet 38 (2019) 70–76.

[28] C. Yin, Y. Ren, A. Adnan, J. Tian, K. Guo, M. Xia, Z. He, D. Zhai, X. Chen, L. Wang, X. Li, X. Qin, S. Li, L. Jin, Title: Developmental validation of Y-SNP pedigree tagging system: A panel via quick ARMS PCR, Forensic Sci Int Genet 46 (2020) 102271.

[29] Z. Zhou, Y. Zhou, Y. Yao, J. Qian, B. Liu, Q. Yang, C. Shao, H. Li, K. Sun, Q. Tang, J. Xie, A 16-plex Y-SNP typing system based on allele-specific PCR for the genotyping of Chinese Y-chromosomal haplogroups, Leg Med (Tokyo) 46 (2020) 101720.

[30] S. Claerhout, P. Verstraete, L. Warnez, S. Vanpaemel, M. Larmuseau, R. Decorte, CSYseq: The first Y-chromosome sequencing tool typing a large number of Y-SNPs and Y-STRs to unravel worldwide human population genetics, PLoS Genet 17(9) (2021) e1009758.

[31] R. Tao, M. Li, S. Chai, R. Xia, Y. Qu, C. Yuan, G. Yang, X. Dong, Y. Bian, S. Zhang, Developmental validation of a 381 Y-chromosome SNP panel for haplogroup analysis in the Chinese populations, Forensic Science International: Genetics 62 (2023) 102803.

[32] M. Wang, G. He, X. Zou, J. Liu, Z. Ye, T. Ming, W. Du, Z. Wang, Y. Hou, Genetic insights into the paternal admixture history of Chinese Mongolians via high-resolution customized Y-SNP SNaPshot panels, Forensic Sci Int Genet 54 (2021) 102565.

[33] M. Lang, H. Liu, F. Song, X. Qiao, Y. Ye, H. Ren, J. Li, J. Huang, M. Xie, S. Chen, M. Song, Y. Zhang, X. Qian, T. Yuan, Z. Wang, Y. Liu, M. Wang, Y. Liu, J. Liu, Y. Hou, Forensic characteristics and genetic analysis of both 27 Y-STRs and 143 Y-SNPs in Eastern Han Chinese population, Forensic Sci Int Genet 42 (2019) e13–e20.

[34] A. Ralf, M. van Oven, D. Montiel Gonzalez, P. de Knijff, K. van der Beek, S. Wootton, R. Lagace, M. Kayser, Forensic Y-SNP analysis beyond SNaPshot: High-resolution Y-chromosomal haplogrouping from low quality and quantity DNA using Ion AmpliSeq and targeted massively parallel sequencing, Forensic Sci Int Genet 41 (2019) 93–106.

[35] T. Gao, L. Yun, D. Zhou, M. Lang, Z. Wang, X. Qian, J. Liu, Y. Hou, Next-generation sequencing of 74 Y-SNPs to construct a concise consensus phylogeny tree for Chinese population, Forensic Science International: Genetics Supplement Series 6 (2017) e96–e98.

[36] J. Liu, L. Jiang, M. Zhao, W. Du, Y. Wen, S. Li, S. Zhang, F. Fang, J. Shen, G. He, Development and validation of a custom panel including 256 Y-SNPs for Chinese Y-chromosomal haplogroups dissection, Forensic Science International: Genetics 61 (2022) 102786.

[37] G. He, M. Wang, X. Zou, H.Y. Yeh, C. Liu, C. Liu, G. Chen, C.C. Wang, Extensive ethnolinguistic diversity at the crossroads of North China and South Siberia reflects multiple sources of genetic diversity, Journal of Systematics and Evolution n/a(n/a) (2022).

[38] G. He, Y. Li, X. Zou, H.Y. Yeh, R. Tang, P. Wang, J. Bai, X. Yang, Z. Wang, J. Guo, J. Chen, J. Chen, M. Yang, J. Zhao, J. Sun, K. Zhu, H. Ma, R. Wang, W. Yang, R. Hu, L.H. Wei, Y. Hou, M. Wang, G. Chen, C.C. Wang, The northern gene flow into southeastern East Asians inferred from genome-wide array genotyping, Journal of Systematics and Evolution n/a(n/a) (2022).

[39] X. Mao, H. Zhang, S. Qiao, Y. Liu, F. Chang, P. Xie, M. Zhang, T. Wang, M. Li, P. Cao, R. Yang, F. Liu, Q. Dai, X. Feng, W. Ping, C. Lei, J.W. Olsen, E.A. Bennett, Q. Fu, The deep population history of northern East Asia from the Late Pleistocene to the Holocene, Cell 184(12) (2021) 3256–3266 e13.

[40] T. Wang, W. Wang, G. Xie, Z. Li, X. Fan, Q. Yang, X. Wu, P. Cao, Y. Liu, R. Yang, F. Liu, Q. Dai, X. Feng, X. Wu, L. Qin, F. Li, W. Ping, L. Zhang, M. Zhang, Y. Liu, X. Chen, D. Zhang, Z. Zhou, Y. Wu, H. Shafiey, X. Gao, D. Curnoe, X. Mao, E.A. Bennett, X. Ji, M.A. Yang, Q. Fu, Human population history at the crossroads of East and Southeast Asia since 11,000 years ago, Cell 184(14) (2021) 3829–3841 e21.

[41] C.C. Wang, H.Y. Yeh, A.N. Popov, H.Q. Zhang, H. Matsumura, K. Sirak, O. Cheronet, A. Kovalev, N. Rohland, A.M. Kim, S. Mallick, R. Bernardos, D. Tumen, J. Zhao, Y.C. Liu, J.Y. Liu, M. Mah, K. Wang, Z. Zhang, N. Adamski, N. Broomandkhoshbacht, K. Callan, F. Candilio, K.S.D. Carlson, B.J. Culleton, L. Eccles, S. Freilich, D. Keating, A.M. Lawson, K. Mandl, M. Michel, J. Oppenheimer, K.T. Ozdogan, K. Stewardson, S. Wen, S. Yan, F. Zalzala, R. Chuang, C.J. Huang, H. Looh, C.C. Shiung, Y.G. Nikitin, A.V. Tabarev, A.A. Tishkin, S. Lin, Z.Y. Sun, X.M. Wu, T.L. Yang, X. Hu, L. Chen, H. Du, J. Bayarsaikhan, E. Mijiddorj, D. Erdenebaatar, T.O. Iderkhangai, E. Myagmar, H. Kanzawa-Kiriyama, M. Nishino, K.I. Shinoda, O.A. Shubina, J. Guo, W. Cai, Q. Deng, L. Kang, D. Li, D. Li, R. Lin Nini, R. Shrestha, L.X. Wang, L. Wei, G. Xie, H. Yao, M. Zhang, G. He, X. Yang, R. Hu, M. Robbeets, S. Schiffels, D.J. Kennett, L. Jin, H. Li, J. Krause, R. Pinhasi, D. Reich, Genomic insights into the formation of human populations in East Asia, Nature 591(7850) (2021) 413–419.

[42] M.A. Yang, X. Fan, B. Sun, C. Chen, J. Lang, Y.C. Ko, C.H. Tsang, H. Chiu, T. Wang, Q. Bao, X. Wu, M. Hajdinjak, A.M. Ko, M. Ding, P. Cao, R. Yang, F. Liu, B. Nickel, Q. Dai, X. Feng, L. Zhang, C. Sun, C. Ning, W. Zeng, Y. Zhao, M. Zhang, X. Gao, Y. Cui, D. Reich, M. Stoneking, Q. Fu, Ancient DNA indicates human population shifts and admixture in northern and southern China, Science 369(6501) (2020) 282–288.

[43] N. Chen, L. Ren, L. Du, J. Hou, V.E. Mullin, D. Wu, X. Zhao, C. Li, J. Huang, X. Qi, M.R. Capodiferro, A. Achilli, C. Lei, F. Chen, B. Su, G. Dong, X. Zhang, Ancient genomes reveal tropical bovid species in the Tibetan Plateau contributed to the prevalence of hunting game until the late Neolithic, Proc Natl Acad Sci U S A 117(45) (2020) 28150–28159.

[44] I. World Medical Association, Declaration of Helsinki. Ethical principles for medical research involving human subjects, Journal of the Indian Medical Association 107(6) (2009) 403–5.

[45] J. Liu, L. Jiang, M. Zhao, W. Du, Y. Wen, S. Li, S. Zhang, F. Fang, J. Shen, G. He, M. Wang, H. Dai, Y. Hou, Z. Wang, Development and validation of a custom panel including 256 Y-SNPs for Chinese Y-chromosomal haplogroups dissection, Forensic Sci Int Genet 61 (2022) 102786.

[46] S. Mallick, H. Li, M. Lipson, I. Mathieson, M. Gymrek, F. Racimo, M. Zhao, N. Chennagiri, S. Nordenfelt, A. Tandon, P. Skoglund, I. Lazaridis, S. Sankararaman, Q. Fu, N. Rohland, G. Renaud, Y. Erlich, T. Willems, C. Gallo, J.P. Spence, Y.S. Song, G. Poletti, F. Balloux, G. van Driem, P. de Knijff, I.G. Romero, A.R. Jha, D.M. Behar, C.M. Bravi, C. Capelli, T. Hervig, A. Moreno-Estrada, O.L. Posukh, E. Balanovska, O. Balanovsky, S. Karachanak-Yankova, H. Sahakyan, D. Toncheva, L. Yepiskoposyan, C. Tyler-Smith, Y. Xue, M.S. Abdullah, A. Ruiz-Linares, C.M. Beall, A. Di Rienzo, C. Jeong, E.B. Starikovskaya, E. Metspalu, J. Parik, R. Villems, B.M. Henn, U. Hodoglugil, R. Mahley, A. Sajantila, G. Stamatoyannopoulos, J.T. Wee, R. Khusainova, E. Khusnutdinova, S. Litvinov, G. Ayodo, D. Comas, M.F. Hammer, T. Kivisild, W. Klitz, C.A. Winkler, D. Labuda, M. Bamshad, L.B. Jorde, S.A. Tishkoff, W.S. Watkins, M. Metspalu, S. Dryomov, R. Sukernik, L. Singh, K. Thangaraj, S. Paabo, J. Kelso, N. Patterson, D. Reich, The Simons Genome Diversity Project: 300 genomes from 142 diverse populations, Nature 538(7624) (2016) 201–206.

[47] L. Pagani, D.J. Lawson, E. Jagoda, A. Morseburg, A. Eriksson, M. Mitt, F. Clemente, G. Hudjashov, M. DeGiorgio, L. Saag, J.D. Wall, A. Cardona, R. Magi, M.A. Wilson Sayres, S. Kaewert, C. Inchley, C.L. Scheib, M. Jarve, M. Karmin, G.S. Jacobs, T. Antao, F.M. Iliescu, A. Kushniarevich, Q. Ayub, C. Tyler-Smith, Y. Xue, B. Yunusbayev, K. Tambets, C.B. Mallick, L. Saag, E. Pocheshkhova, G. Andriadze, C. Muller, M.C. Westaway, D.M. Lambert, G. Zoraqi, S. Turdikulova, D. Dalimova, Z. Sabitov, G.N.N. Sultana, J. Lachance, S. Tishkoff, K. Momynaliev, J. Isakova, L.D. Damba, M. Gubina, P. Nymadawa, I. Evseeva, L. Atramentova, O. Utevska, F.X. Ricaut, N. Brucato, H. Sudoyo, T. Letellier, M.P. Cox, N.A. Barashkov, V. Skaro, L. Mulahasanovic, D. Primorac, H. Sahakyan, M. Mormina, C.A. Eichstaedt, D.V. Lichman, S. Abdullah, G. Chaubey, J.T.S. Wee, E. Mihailov, A. Karunas, S. Litvinov, R. Khusainova, N. Ekomasova, V. Akhmetova, I. Khidiyatova, D. Marjanovic, L. Yepiskoposyan, D.M. Behar, E. Balanovska, A. Metspalu, M. Derenko, B. Malyarchuk, M. Voevoda, S.A. Fedorova, L.P. Osipova, M.M. Lahr, P. Gerbault, M. Leavesley, A.B. Migliano, M. Petraglia, O. Balanovsky, E.K. Khusnutdinova, E. Metspalu, M.G. Thomas, A. Manica, R. Nielsen, R. Villems, E. Willerslev, T. Kivisild, M. Metspalu, Genomic analyses inform on migration events during the peopling of Eurasia, Nature 538(7624) (2016) 238–242.

[48] C.C. Chang, C.C. Chow, L.C. Tellier, S. Vattikuti, S.M. Purcell, J.J. Lee, Second-generation PLINK: rising to the challenge of larger and richer datasets, Gigascience 4 (2015) 7.

[49] A. Jagadeesan, S.S. Ebenesersdottir, V.B. Guethmundsdottir, E.L. Thordardottir, K.H.S. Moore, A. Helgason, HaploGrouper: a generalized approach to haplogroup classification, Bioinformatics 37(4) (2021) 570–572.

[50] H. Chen, Y. Lu, D. Lu, S. Xu, Y-LineageTracker: a high-throughput analysis framework for Y-chromosomal next-generation sequencing data, BMC Bioinformatics 22(1) (2021) 114.

[51] S. Dellicour, M.S. Gill, N.R. Faria, A. Rambaut, O.G. Pybus, M.A. Suchard, P. Lemey Relax, Keep Walking — A Practical Guide to Continuous Phylogeographic Inference with BEAST, Molecular Biology and Evolution 38(8) (2021) 3486–3493.

[52] J.W. Leigh, D. Bryant, popart: full-feature software for haplotype network construction, Methods in Ecology and Evolution 6(9) (2015) 1110–1116.

[53] J. Chen, G. He, Z. Ren, Q. Wang, Y. Liu, H. Zhang, M. Yang, H. Zhang, J. Ji, J. Zhao, J. Guo, K. Zhu, X. Yang, R. Wang, H. Ma, C.C. Wang, J. Huang, Genomic Insights Into the Admixture History of Mongolic- and Tungusic-Speaking Populations From Southwestern East Asia, Front Genet 12(880) (2021) 685285.

[54] Q. Wu, H.Z. Cheng, N. Sun, P.C. Ma, J. Sun, H.B. Yao, Y.M. Xie, Y.L. Li, S.L. Meng, M. Zhabagin, Y.H. Cai, D.R. Lu, S. Yan, L.H. Wei, Phylogenetic analysis of the Y-chromosome haplogroup C2b-F1067, a dominant paternal lineage in Eastern Eurasia, J Hum Genet 65(10) (2020) 823–829.

[55] P. Zhang, H. Luo, Y. Li, Y. Wang, J. Wang, Y. Zheng, Y. Niu, Y. Shi, H. Zhou, T. Song, Q. Kang, K.I. Han, T. Xu, S. He, NyuWa Genome resource: A deep whole-genome sequencing-based variation profile and reference panel for the Chinese population, Cell Rep 37(7) (2021) 110017.

[56] Y. Cao, L. Li, M. Xu, Z. Feng, X. Sun, J. Lu, Y. Xu, P. Du, T. Wang, R. Hu, Z. Ye, L. Shi, X. Tang, L. Yan, Z. Gao, G. Chen, Y. Zhang, L. Chen, G. Ning, Y. Bi, W. Wang, M.A.P.C. China, The ChinaMAP analytics of deep whole genome sequences in 10,588 individuals, Cell Res 30(9) (2020) 717–731.

[57] G.L. He, M.G. Wang, Y.X. Li, X. Zou, H.Y. Yeh, R.K. Tang, X.M. Yang, Z. Wang, J.X. Guo, T. Luo, J. Zhao, J. Sun, R. Hu, L.H. Wei, G. Chen, Y.P. Hou, C.C. Wang, Fine-scale north-to-south genetic admixture profile in Shaanxi Han Chinese revealed by genome - wide demographic history reconstruction, Journal of Systematics and Evolution (2021) 1–20.

[58] M. Xie, F. Song, J. Li, M. Lang, H. Luo, Z. Wang, J. Wu, C. Li, C. Tian, W. Wang, H. Ma, Z. Song, Y. Fan, Y. Hou, Genetic substructure and forensic characteristics of Chinese Hui populations using 157 Y-SNPs and 27 Y-STRs, Forensic Science International: Genetics 41 (2019) 11–18.

[59] F. Wang, F. Song, M. Song, H. Luo, Y. Hou, Genetic structure and paternal admixture of the modern Chinese Zhuang population based on 37 Y-STRs and 233 Y-SNPs, Forensic Science International: Genetics 58 (2022) 102681.

[60] R. Tao, M. Li, S. Chai, R. Xia, Y. Qu, C. Yuan, G. Yang, X. Dong, Y. Bian, S. Zhang, C. Li, Developmental validation of a 381 Y-chromosome SNP panel for haplogroup analysis in the Chinese populations, Forensic Science International: Genetics 62 (2023) 102803.

[61] M. Song, Z. Wang, Y. Zhang, C. Zhao, M. Lang, M. Xie, X. Qian, M. Wang, Y. Hou, Forensic characteristics and phylogenetic analysis of both Y-STR and Y-SNP in the Li and Han ethnic groups from Hainan Island of China, Forensic Science International: Genetics 39 (2019) e14–e20.

[62] M. Song, Z. Wang, Q. Lyu, J. Ying, Q. Wu, L. Jiang, F. Wang, Y. Zhou, F. Song, H. Luo, Y. Hou, X. Song, B. Ying, Paternal genetic structure of the Qiang ethnic group in China revealed by high-resolution Y-chromosome STRs and SNPs, Forensic Science International: Genetics 61 (2022) 102774.

[63] Y.Z. Huang, L.H. Wei, S. Yan, S.Q. Wen, C.C. Wang, Y.J. Yang, L.X. Wang, Y. Lu, C. Zhang, S.H. Xu, D.L. Yao, L. Jin, H. Li, Whole sequence analysis indicates a recent southern origin of Mongolian Y-chromosome C2c1a1a1-M407, Mol Genet Genomics 293(3) (2018) 657–663.

[64] H. Chen, R. Lin, Y. Lu, R. Zhang, Y. Gao, Y. He, S. Xu, Tracing Bai-Yue ancestry in aboriginal Li people on Hainan Island, Mol Biol Evol (2022) msac210.

[65] G. He, Z. Wang, J. Guo, M. Wang, X. Zou, R. Tang, J. Liu, H. Zhang, Y. Li, R. Hu, L.H. Wei, G. Chen, C.C. Wang, Y. Hou, Inferring the population history of Tai-Kadai-speaking people and southernmost Han Chinese on Hainan Island by genome-wide array genotyping, Eur J Hum Genet 28(8) (2020) 1111–1123.

[66] G. He, Z. Wang, M. Wang, T. Luo, J. Liu, Y. Zhou, B. Gao, Y. Hou, Forensic ancestry analysis in two Chinese minority populations using massively parallel sequencing of 165 ancestry-informative SNPs, Electrophoresis 39(21) (2018) 2732–2742.

[67] X. Ma, W. Yang, Y. Gao, Y. Pan, Y. Lu, H. Chen, D. Lu, S. Xu, Genetic Origins and Sex-Biased Admixture of the Huis, Mol Biol Evol 38(9) (2021) 3804–3819.

[68] G. He, Z.-Q. Fan, X. Zou, X. Deng, H.-Y. Yeh, Z. Wang, J. Liu, Q. Xu, L. Chen, X.-H. Deng, C.-C. Wang, C. Liu, M. Wang, C. Liu, Demographic model and biological adaptation inferred from the genome-wide single nucleotide polymorphism data reveal tripartite origins of southernmost Chinese Huis, American Journal of Biological Anthropology n/a(n/a).

[69] C. Jeong, A.T. Ozga, D.B. Witonsky, H. Malmstrom, H. Edlund, C.A. Hofman, R.W. Hagan, M. Jakobsson, C.M. Lewis, M.S. Aldenderfer, A. Di Rienzo, C. Warinner, Long-term genetic stability and a high-altitude East Asian origin for the peoples of the high valleys of the Himalayan arc, Proc Natl Acad Sci U S A 113(27) (2016) 7485–90.

[70] G. He, M. Wang, X. Zou, P. Chen, Z. Wang, Y. Liu, H. Yao, L.H. Wei, R. Tang, C.C. Wang, H.Y. Yeh, Peopling History of the Tibetan Plateau and Multiple Waves of Admixture of Tibetans Inferred From Both Ancient and Modern Genome-Wide Data, Front Genet 12(1634) (2021) 725243.

[71] Y. Liu, J. Xie, M. Wang, C. Liu, J. Zhu, X. Zou, W. Li, L. Wang, C. Leng, Q. Xu, H.Y. Yeh, C.C. Wang, X. Wen, C. Liu, G. He, Genomic Insights Into the Population History and Biological Adaptation of Southwestern Chinese Hmong-Mien People, Front Genet 12 (2021) 815160.

[72] L. Li, P. Huang, X. Sun, S. Wang, M. Xu, S. Liu, Z. Feng, Q. Zhang, X. Wang, X. Zheng, M. Dai, Y. Bi, G. Ning, Y. Cao, W. Wang, The ChinaMAP reference panel for the accurate genotype imputation in Chinese populations, Cell Res 31(12) (2021) 1308–1310.

